# No evidence of SARS-CoV-2 infection in Neotropical Primates sampled during COVID-19 pandemic in Minas Gerais and Rio Grande do Sul, Brazil

**DOI:** 10.1101/2021.06.17.448890

**Authors:** Filipe Vieira Santos de Abreu, Mariana Viana Macedo, Alex Junio Jardim da Silva, Cirilo Henrique de Oliveira, Vinícius de Oliveira Ottone, Marco Antônio Barreto de Almeida, Edmilson dos Santos, Jader da Cruz Cardoso, Aline Scarpellini Campos, Claudia Maria Dornelles da Silva, Amanda Gonzales da Silva, Miguel de Souza Andrade, Valéria Magro Octaviano Bernis, Walter Octaviano Bernis Filho, Giliane de Souza Trindade, George Rego Albuquerque, Anaiá da Paixão Sevá, Bergmann Morais Ribeiro, Danilo Simonini Teixeira, Fabrício Souza Campos, Ana Cláudia Franco, Paulo Michel Roehe, Danilo Bretas de Oliveira

## Abstract

In 2019, a new coronavirus disease (COVID-19) was detected in China. Severe acute respiratory syndrome coronavirus 2 (SARS-CoV-2) was capable to infect domestic and captive mammals like cats, tigers and minks. Due to genetic similarities, concern about the infection of Non-Human Primates (NHPs) and the establishment of a sylvatic cycle has grown in the Americas. In this study, neotropical primates (NP) were sampled in different areas from Brazil to investigate whether they were infected by SARS-CoV-2. A total of 89 samples from 51 NP of four species were examined. No positive samples were detected via RT-qPCR, regardless of the NHP species, tissue or habitat tested. This work provides the first report on the lack of evidence of circulation of SARS-CoV-2 in NP. The expand of wild animals sampling is necessary to understand their role in the epidemiology of SARS-CoV-2.

---

In December 2019, a new disease caused by a virus belonging to *Coronaviridae* family, was first detected in the city of Wuhan, China, and was named coronavirus disease (COVID-19) (Wu et al. 2020). The etiologic agent - Severe Acute Respiratory Syndrome Coronavirus 2 (SARS-CoV-2) - is highly transmissible among humans through droplets of saliva and quickly spread across the planet. In March 2020, the virus reached every continent which led the World Health Organization to declare COVID-19 as a pandemic (World Health Organization).

Initial studies of epidemiology and genetic comparison showed that the new virus has high similarity to other coronaviruses found in bats and pangolin (96 and 91% similarity, respectively) (Zhang et al. 2020) and that the “spillover”, that is, the “jump” between hosts could have occurred in the vicinity of the popular market in Wuhan, where these animals are traditionally traded for human consumption (Wu et al. 2020). Thenceforth, it has been questioned whether SARS-CoV-2 would be able to perform a “spillback” - transmission from humans to wild animals -, which could generate wild cycles of the virus, making it even more difficult to face.

In this regard, concern about the infection of non-human primates (NHPs) has grown since the biological, genetic and biochemical similarity, could facilitate the transmission of SARS-CoV-2 to NHPs. Comparative studies have indicated that the cellular receptor angiotensin-converting enzyme 2 (ACE2), used by this virus for adsorption to cells, is similar between humans and NHPs, especially in Old World species; as such, these have been used as experimental models for the study of SARS-CoV-2 (Melin et al. 2020). In the Americas, there is a great concern about the risks of SARS-CoV-2 spillback from humans to wildlife, including neotropical NHPs. This could have at least two impacts: a) establishment of a sylvatic cycle with serious implications on control or eradication efforts, and: b) unforeseeable impact on biodiversity, especially if endangered NHP species become involved. The historical comparison with yellow fever virus (YFV), whose sylvatic cycle was first recognized almost a century ago after a spillback from human to NHPs, exemplifies the potential damage that may be associated to such impacts (Possas et al. 2018).

To our knowledge, the only neotropical primate species experimentally infected with SARS-CoV-2 was the marmoset *Callithrix jacchus*. Infected individuals had pyrexia and viral RNA loads detectable for several days post-infection in oral, nasal and anal swabs, as well as in blood and stool samples (Lu et al. 2020). However, little is known about the possibility of viral infection in natural conditions and in other neotropical primates’ families or species. Worryingly, in many Brazilian cities where cases of COVID-19 are currently reaching unprecedented levels, *Callithrix* spp. are adapted to synanthropic environments, often kept as pets and feeding on human foods. Such coexistence might represent a significant potential source of infection (Longa et al. 2011). In addition, the rapid dissemination of SARS-CoV-2 currently verified throughout Brazil can lead to the exposure of other NHPs, including wild species.

The present study was carried out in search for serological and virological evidences of SARS-CoV-2 infections in NHPs from urban, sylvatic and rural areas of two Brazilian states, Minas Gerais and Rio Grande do Sul. The current study is part of a wider NHP sampling effort whose main goal is to contribute to the national program for epidemiological surveillance of yellow fever virus.

Neotropical Primates’ samplings were performed before (November 2019) and during (July 2020 - February 2021) the COVID-19 pandemic in two Brazilian states. First in Minas Gerais (Southeast region), where the savannah-like biome predominates, captures occurred in four (Fig. 1A). After in Rio Grande do Sul (South region), covered by Atlantic Forest and Pampa biomes, in seven municipalities (Fig. 1B). Human COVID-19 cases per 100.000 inhabitants up to the sampling efforts are highlighted by the colors of the municipalities on the map (SES-MG 2020; SES-RS 2021).

**Figure 1:**
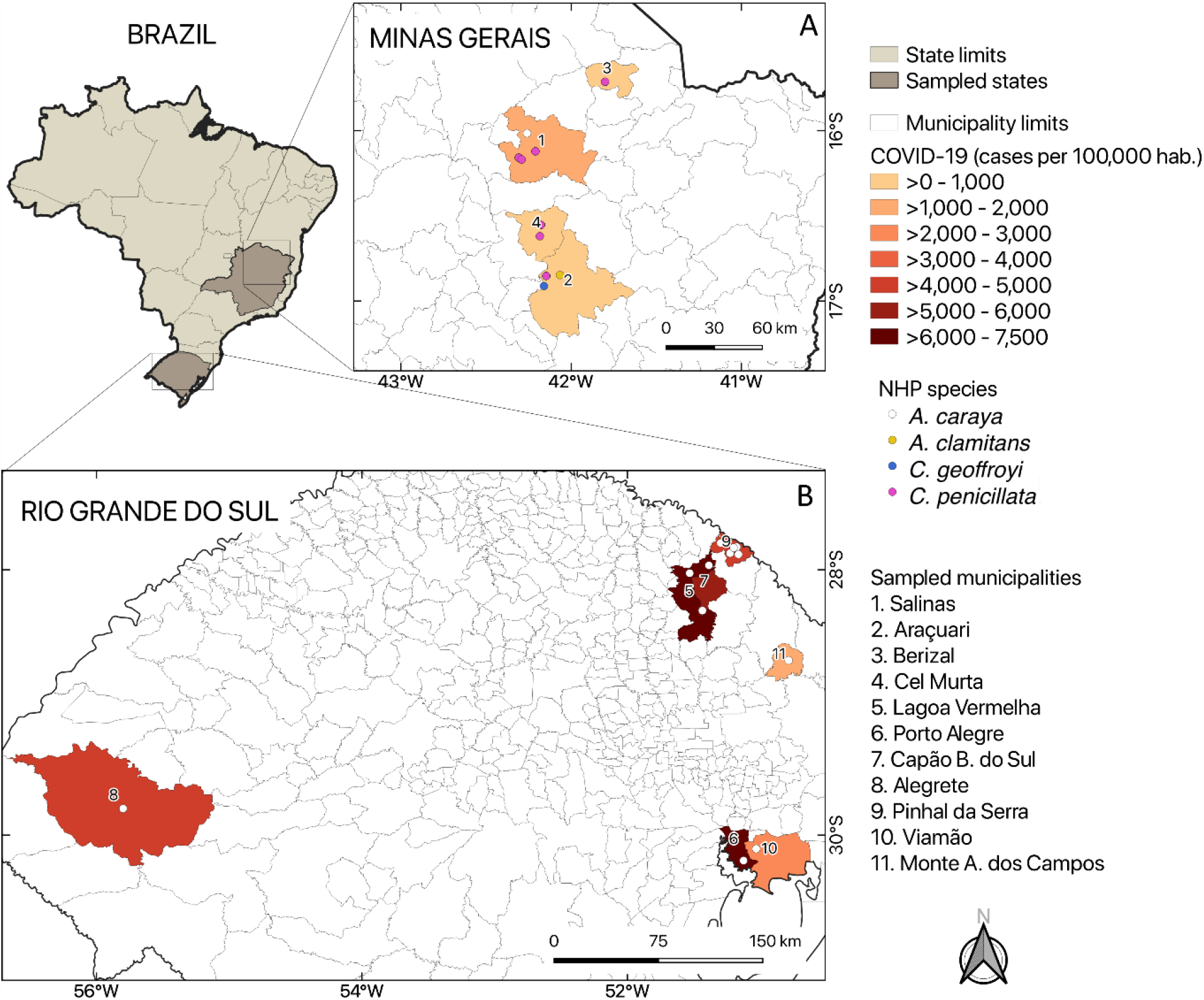
Municipalities where sampling of NHPs was performed. A: Minas Gerais state; B: Rio Grande do Sul state. The number of COVID-19 human cases in each municipality were counted until the sample collection date. Data were obtained from state health departments (SES-MG 2020; SES-RS 2021). The figure was done using QGIS software, version 3.10.

Live marmosets were captured with Tomahawk traps, according to the protocol previously described (Abreu et al. 2019a). Dead or sick marmosets and howler monkeys were also evaluated. In MG, they were found based on an information network made up of health professionals, farmers, and other people in contact with natural environments, as described elsewhere (Abreu et al. 2019b) while in RS, Health Department network people found and informed the epizooties. Serum and/or oral swab, and viscera (lung, liver, spleen or kidney of dead animals) were collected and preserved in RNA*later* (Thermo Fisher™) and stored in liquid nitrogen at - 186°C. Our methods and protocols were previously approved by the Institutional Ethics Committee for Animal Experimentation (Protocol CEUA/IFNMG n°14/2019) and by Brazilian Ministry of the Environment (SISBIO n° 71714-2).

RNA extractions were performed using QIAamp Viral RNA Mini Kit (QIAGEN® (for serum) and TRIzol™ Reagent (Thermo Fisher™) (for viscera) following manufacturer’s instructions. Detection of the SARS-CoV-2 viral RNA in samples from Minas Gerais was performed using a real-time RT-PCR kit (CDC 2019-nCoV Real-Time RT-PCR Diagnostic Panel, which targets the N1 (2019-nCoV_N1 Combined Primer/Probe Mix) and N2 (2019-nCoV_N2 Combined Primer/Probe Mix) genes of the SARS-CoV-2 (CDC Division of Viral Diseases 2020). In Rio Grande do Sul, detection was performed using the Allplex™ 2019-nCoV (Seegene®), which targets genomic regions within the *E, RdRP*, and *N* genes.

In total, 51 NHPs belonging to two different families (Callitrichidae and Atelidae) and four species (*Callithrix penicillata, C. jacchus, Alouatta caraya* and *A. guariba clamitans*) were examined (Table 1). Seven NHPs were captured before the reported virus introduction (august - november / 2019) and 44 after the introduction (july 2020 to february 2021). Different tissues were analyzed, totalizing 89 samples: 30 of serum, 20 of oral swabs, 19 of liver, 14 of kidney, 3 of lung and 3 of whole blood (Table 1). No positive samples were detected, regardless of NHP species, tissue, habitat, municipalities or state/biome tested (Table 1).

**Table 1:**
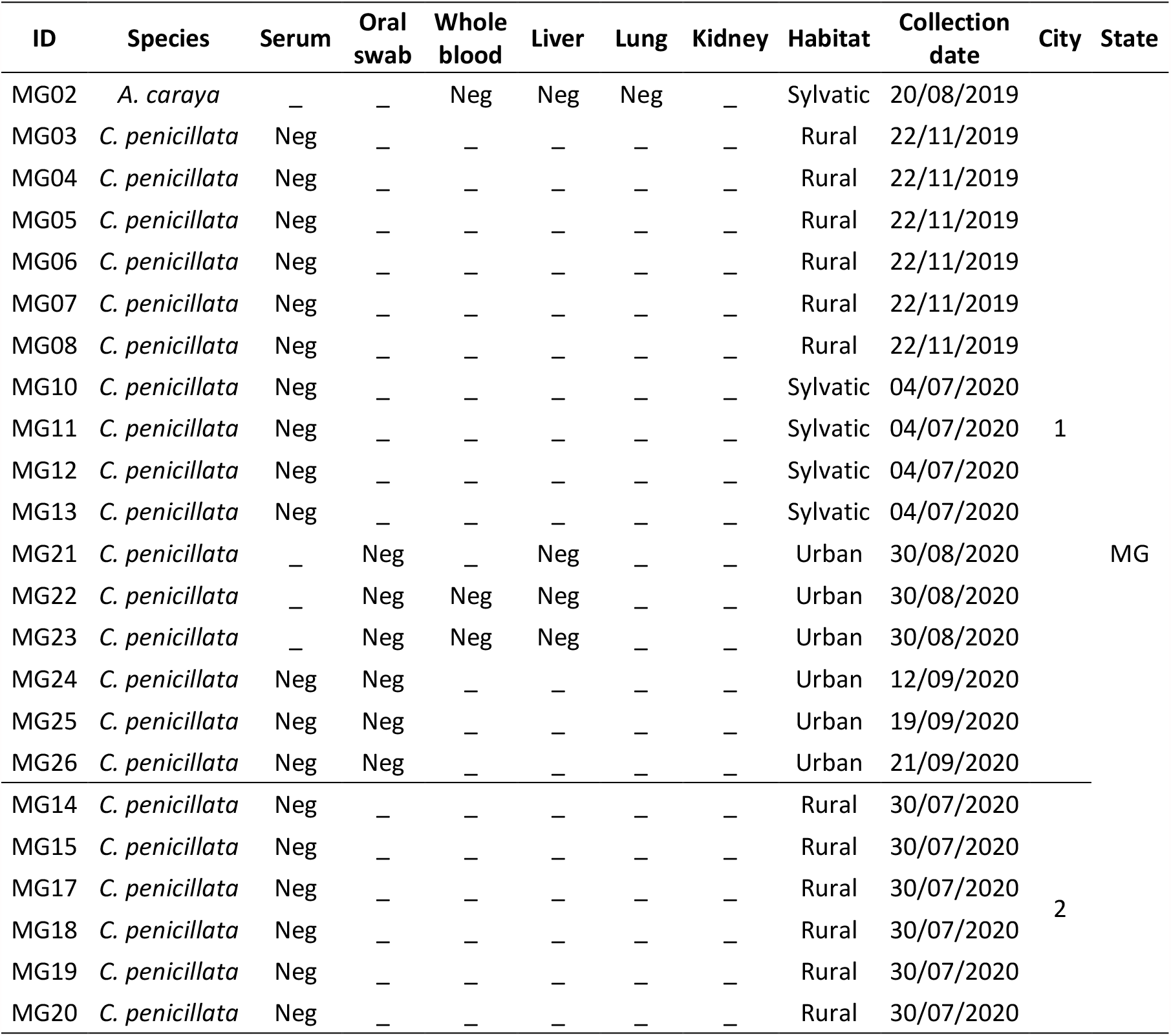
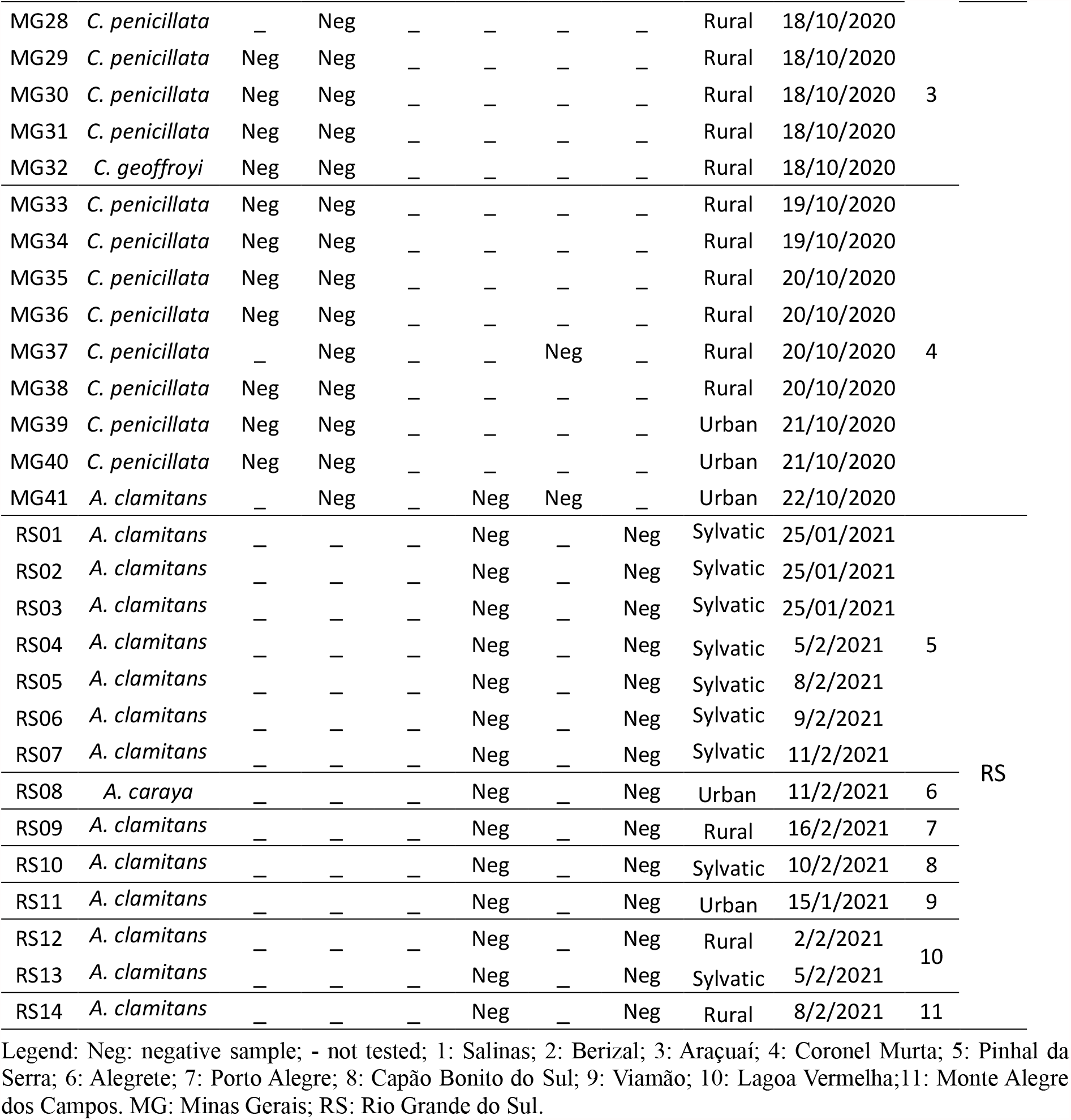
Description of samples tested by species, tissue, habitat, date, municipality (city) and state.

This is the first study evaluating the natural infection of free-living neotropical NHPs with SARS-CoV-2. We found no evidence of SARS-CoV-2 infection in NHPs from urban, rural or sylvatic habitats regardless of the tissue or species surveyed. Even so, it is important to maintain and expand the surveillance of wildlife, aiming at the early detection of spillover / spillback among human beings and other animals, considering the widespread of SARS-CoV-2 in Brazil.

Since the emergence, reports of natural SARS-CoV-2 infection of several animals have accumulated, including domestic animals like cats (Sailleau et al. 2020; Carlos et al. 2021) and wild captive animals like tigers (McAloose et al. 2020) and minks (Hammer et al. 2021). Minks have also been described as a source of infection for humans (Hammer et al. 2021). In addition, experimental infection studies have reported susceptibility of several animals such as ferrets, bat, hamsters (Shi et al. 2020; Imai et al. 2020), and mainly Old World NHPs (Zheng et al. 2020; Lu et al. 2020). As a result, concern about the SARS-CoV-2 spillback for Neotropical Primates – as happened with YFV in the last century and probably with ZIKA in the last decade – has grown (Terzian et al. 2018; Yunes Guimarães et al. 2020).

The only report of experimental infection in a neotropical NHP published to date, examined the dynamics of SARS-CoV-2 infection in *Callithrix jacchus*, the common marmoset. Viral genomes were detected in oral, nasal and anal swabs as well as in serum and feces, despite the lower susceptibility of the species when compared to Old World Primates. (Lu et al. 2020). In the present study, all tissues sampled from the 44 free-living primates collected after the introduction of the virus in Brazil were negative, including samples from NHP living in rural and urban environments, in close contact with humans.

The cellular receptor ACE2 used by the virus for adsorption to cells, is a major determinant in the susceptibility of animals to SARS-CoV-2 infection. The ACE2 receptor of cells from Old World NHPs have up to 99% identity with human ACE2 protein, what might contribute significantly to the high susceptibility that such species display to infection with this virus (Shi et al. 2020; Melin et al. 2020). Neotropical NHPs, such as those sampled in the present study, have ACE2 receptors that share about 92% identity to human ACE2, with 4 differences across the twelve binding sites regarded as essential for the ACE2 protein to allow binding of SARS-CoV-2 (Melin et al. 2020; Joyraj Bhattacharjee et al. 2021). This may be related to species’ differences in susceptibility and, possibly, with our negative results, even in NHPs living in urban areas with high SARS-CoV-2 human incidence. Even so, it is important to investigate whether the new variants of SARS-CoV-2 could increase their infectivity in wild animals.

Brazil is the country with the greatest biodiversity on the planet and concentrates the largest number of NHP species, many of these endangered due to anthropogenic effects such as deforestation which, in addition to drastically reducing habitats, increases the risks of zoonotic transmissions (Paglia et al. 2012; Guthid et al. 2020). The explosive spread of SARS-CoV-2 in the country and the close contact between humans and native fauna, provides conditions for spillovers and spillback which increases the need for surveillance for early identification and management of potential harmful effects (Gryseels et al. 2020). In this sense and despite the small sampling, our study contributes to shedding light on the subject by providing the first results about the absence of circulation of SARS-CoV-2 in Neotropical Primates. It is urgent to expand the sampling of NHP living in different environments, to include other primate species and even other mammals to carry out an active surveillance of wild animals and understand the epidemiology of SARS-CoV-2 in natural environments.

## Acknowledgement

The authors thanks to all institutions and technicians that contributed and supported the field work. We are especially grateful to Centro de Estudo Multidisciplinar em Conservação Animal IFNMG – Salinas (CEMCA), Aline Tátila Ferreira, Sandy Micaele, Maria Eduarda and Pedro Augusto, for their valuable contribution during the field work. We are also grateful to members of Secretaria de Saúde and Centro de Controle de Zoonoses de Salinas, Berizal, Rubelita, and Coronel Murta; and Rio Grande do Sul State Health Department: Members of Yellow Fever Surveillance State Reference Team. We also thanks to Conselho Nacional de Desenvolvimento Científico e Tecnológico (CNPq) for scholarship awarded.

## Ethics approval

All applicable institutional and/or national guidelines for the care and use of animals were followed. Methods and protocols were previously approved by the Institutional Ethics Committee for Animal Experimentation (Protocol CEUA/IFNMG n°14/2019) and by Brazilian Ministry of the Environment (SISBIO n° 71714-2). This article does not contain any studies with human participants.

## References

Abreu FVS de, Santos E dos, Mello ARL, Gomes LR, Alvarenga DAM de, Gomes MQ, Vargas WP, Bianco-Júnior C, Pina-Costa A de, Teixeira DS, Romano APM, Manso PP de A, Pelajo-Machado M, Brasil P, Daniel-Ribeiro CT, Brito CFA de, Ferreira-da-Cruz M de F, Lourenço-de-Oliveira R (2019a) Howler monkeys are the reservoir of malarial parasites causing zoonotic infections in the Atlantic forest of Rio de Janeiro. PLoS Negl Trop Dis 13:e0007906. https://doi.org/10.1371/journal.pntd.0007906

Abreu FVS, dos Santos E, Gomes MQ, Vargas WP, Oliveira Passos PH, Nunes e Silva C, Araújo PC, Pires JR, Romano APM, Teixeira DS, Lourenço-de-Oliveira R (2019b) Capture of Alouatta guariba clamitans for the surveillance of sylvatic yellow fever and zoonotic malaria: Which is the best strategy in the tropical Atlantic Forest? Am J Primatol 81:e23000. https://doi.org/10.1002/ajp.23000

Carlos RSA, Mariano APM, Maciel BM, Gadelha SR, Melo Silva M, Belitardo EMMA, Rocha Djpg, Almeida JPP, Pacheco LGC, Aguiar Ergr, Fehlberg HF, Albuquerque GR (2021) First genome sequencing of SARS-CoV-2 recovered from an infected cat and its owner in Latin America. Transbound Emerg Dis tbed.13984. https://doi.org/10.1111/tbed.13984

CDC Division of Viral Diseases (2020) CDC 2019-Novel Coronavirus (2019-nCoV) Real-Time RT-PCR Diagnostic Panel 2019-nCoVEUA-01 CDC-006-00019, Revision: 06. https://www.fda.gov/media/134922/download. Accessed 5 Dec 2020

Gryseels S, De Bruyn L, Gyselings R, Calvignac-Spencer S, Leendertz FH, Leirs H (2020) Risk of human-to-wildlife transmission of SARS-CoV-2. Mamm Rev. https://doi.org/10.1111/mam.12225

Guthid S, Hanley KA, Althouse BM, Boots M (2020) Ecological processes underlying the emergence of novel enzootic cycles: Arboviruses in the neotropics as a case study. PLoS Negl. Trop. Dis. 14:1–22

Hammer AS, Quaade ML, Rasmussen TB, Fonager J, Rasmussen M, Mundbjerg K, Lohse L, Strandbygaard B, Jørgensen CS, Alfaro-Núñez A, Rosenstierne MW, Boklund A, Halasa T, Fomsgaard A, Belsham GJ, Bøtner A (2021) SARS-CoV-2 transmission between mink (neovison vison) and Humans, Denmark. Emerg Infect Dis 27:547–551. https://doi.org/10.3201/eid2702.203794

Imai M, Iwatsuki-Horimoto K, Hatta M, Loeber S, Halfmann PJ, Nakajima N, Watanabe T, Ujie M, Takahashi K, Ito M, Yamada S, Fan S, Chiba S, Kuroda M, Guan L, Takada K, Armbrust T, Balogh A, Furusawa Y, Okuda M, Ueki H, Yasuhara A, Sakai-Tagawa Y, Lopes TJS, Kiso M, Yamayoshi S, Kinoshita N, Ohmagari N, Hattori SI, Takeda M, Mitsuya H, Krammer F, Suzuki T, Kawaoka Y (2020) Syrian hamsters as a small animal model for SARS-CoV-2 infection and countermeasure development. Proc Natl Acad Sci U S A 117:16587–16595. https://doi.org/10.1073/pnas.2009799117

Joyraj Bhattacharjee M, Lin J-J, Chang C-Y, Chiou Y-T, Li T-N, Tai C-W, Shiu T-F, Chen C-A, Chou C-Y, Chakraborty P, Yuan Tseng Y, Hui-Ching Wang L, Li W-H (2021) Identifying primate ACE2 variants that confer resistance to SARS-CoV-2. Mol Biol Evol. https://doi.org/10.1093/molbev/msab060

Longa CS, Bruno SF, Pires AR, Romijn PC, Kimura LS, Costa CHC (2011) Human herpesvirus 1 in wild marmosets, Brazil, 2008. Emerg. Infect. Dis. 17:1308–1310

Lu S, Zhao Y, Yu W, Yang Y, Gao J, Wang J, Kuang D, Yang M, Yang J, Ma C, Xu J, Qian X, Li H, Zhao S, Li J, Wang H, Long H, Zhou J, Luo F, Ding K, Wu D, Zhang Y, Dong Y, Liu Y, Zheng Y, Lin X, Jiao L, Zheng H, Dai Q, Sun Q, Hu Y, Ke C, Liu H, Peng X (2020) Comparison of nonhuman primates identified the suitable model for COVID-19. Signal Transduct Target Ther 5:1–9. https://doi.org/10.1038/s41392-020-00269-6

McAloose D, Laverack M, Wang L, Killian ML, Caserta LC, Yuan F, Mitchell PK, Queen K, Mauldin MR, Cronk BD, Bartlett SL, Sykes JM, Zec S, Stokol T, Ingerman K, Delaney MA, Fredrickson R, Ivančić M, Jenkins-Moore M, Mozingo K, Franzen K, Bergeson NH, Goodman L, Wang H, Fang Y, Olmstead C, McCann C, Thomas P, Goodrich E, Elvinger F, Smith DC, Tong S, Slavinski S, Calle PP, Terio K, Torchetti MK, Diel DG (2020) From people to panthera: Natural sars-cov-2 infection in tigers and lions at the bronx zoo. MBio 11:1–13. https://doi.org/10.1128/mBio.02220-20

Melin AD, Janiak MC, Marrone F, Arora PS, Higham JP (2020) Comparative ACE2 variation and primate COVID-19 risk. Commun Biol 3:1–9. https://doi.org/10.1038/s42003-020-01370-w

Paglia AP, Rylands AB, Herrmann G, Aguiar LMS, Chiarello AG, Leite YLR, Costa LP, Siciliano S (2012) Lista anotada dos mamiferos do Brasil, segunda edicao, 2nd edn. Conservation International, Arlington

Possas C, Lourenço-de-oliveira R, Tauil PL, Pinheiro FDP, Pissinatti A, Venâncio R, Freire M, Martins RM, Homma A (2018) Yellow fever outbreak in Brazil : the puzzle of rapid viral spread and challenges for immunisation. Mem Inst Oswaldo Cruz 113:1–12. https://doi.org/10.1590/0074-02760180278

Sailleau C, Dumarest M, Vanhomwegen J, Delaplace M, Caro V, Kwasiborski A, Hourdel V, Chevaillier P, Barbarino A, Comtet L, Pourquier P, Klonjkowski B, Manuguerra J, Zientara S, Le Poder S (2020) First detection and genome sequencing of SARS-CoV-2 in an infected cat in France. Transbound Emerg Dis 67:2324–2328. https://doi.org/10.1111/tbed.13659

SES-MG (2020) COVID-19 Coronavírus Boletim Minas Gerais. Belo Horizonte

SES-RS (2021) SES/RS - Coronavirus. https://ti.saude.rs.gov.br/covid19/. Accessed 22 Mar 2021

Shi J, Wen Z, Zhong G, Yang H, Wang C, Huang B, Liu R, He X, Shuai L, Sun Z, Zhao Y, Liu P, Liang L, Cui P, Wang J, Zhang X, Guan Y, Tan W, Wu G, Chen H, Bu Z (2020) Susceptibility of ferrets, cats, dogs, and other domesticated animals to SARS–coronavirus 2. Science (80-) 7015:eabb7015. https://doi.org/10.1126/science.abb7015

Terzian ACB, Zini N, Sacchetto L, Rocha RF, Parra MCP, Del Sarto JL, Dias ACF, Coutinho F, Rayra J, da Silva RA, Costa VV, Fernandes NCCDA, Réssio R, Díaz-Delgado J, Guerra J, Cunha MS, Catão-Dias JL, Bittar C, Reis AFN, Santos INP dos, Ferreira ACM, Cruz Leaa, Rahal P, Ullmann L, Malossi C, Araújo Jr JP de, Widen S, de Rezende IM, Mello É, Pacca CC, Kroon EG, Trindade G, Drumond B, Chiaravalloti-Neto F, Vasilakis N, Teixeira MM, Nogueira ML (2018) Evidence of natural Zika virus infection in neotropical non-human primates in Brazil. Sci Rep 8:16034. https://doi.org/10.1038/s41598-018-34423-6

World Health Organization WHO Director-General’s opening remarks at the media briefing on COVID-19 - 11 March 2020. https://www.who.int/director-general/speeches/detail/who-director-general-s-opening-remarks-at-the-media-briefing-on-covid-1911-march-2020. Accessed 23 Mar 2021

Wu F, Zhao S, Yu B, Chen YM, Wang W, Song ZG, Hu Y, Tao ZW, Tian JH, Pei YY, Yuan ML, Zhang YL, Dai FH, Liu Y, Wang QM, Zheng JJ, Xu L, Holmes EC, Zhang YZ (2020) A new coronavirus associated with human respiratory disease in China. Nature 579:265–269. https://doi.org/10.1038/s41586-020-2008-3

Yunes Guimarães V, Augusto Justo A, Luís Martins L, Catão-Dias Jl, Sacristán C (2020) EMERGING CORONAVIRUSES IN NEOTROPICAL PRIMATES: A NEW THREAT? Rev Ciência Veterinária e Saúde Pública 7:001–012. https://doi.org/10.4025/revcivet.v7i1.55490

Zhang T, Wu Q, Zhang Z (2020) Probable Pangolin Origin of SARS-CoV-2 Associated with the COVID-19 Outbreak. Curr Biol 30:1346-1351.e2. https://doi.org/10.1016/j.cub.2020.03.022

Zheng H, Li H, Guo L, Liang Y, Li J, Wang X, Hu Y, Wang L, Liao Y, Yang F, Li Y, Fan S, Li D, Cui P, Wang Q, Shi H, Chen Y, Yang Z, Yang J, Shen D, Cun W, Zhou X, Dong X, Wang Y, Chen Y, Dai Q, Jin W, He Z, Li Q, Liu L (2020) Virulence and pathogenesis of SARS-CoV-2 infection in rhesus macaques: A nonhuman primate model of COVID-19 progression. PLoS Pathog 16:e1008949. https://doi.org/10.1371/JOURNAL.PPAT.1008949

